# MTDH-SND1 disruption sensitizes ovarian cancer to ferroptosis and PARP inhibition

**DOI:** 10.64898/2026.05.18.725896

**Authors:** Parisa Esmaeili, Ahmad Nasimian, Elsa Ernestål, Elna Persson, Bianca Bochis, Yuan Li, Måns Zamore, Anna Sandström Gerdtsson, Julhash U Kazi, Fredrik Levander

**Affiliations:** Department of Immunotechnology, Lund University, Lund, Sweden; Division of Translational Cancer Research, Department of Laboratory Medicine, Lund University, Lund, Sweden; Department of Immunotechnology, National Bioinformatics Infrastructure Sweden, Science for Life laboratory, Lund University, Lund, Sweden; Lund Stem Cell Center, Department of Laboratory Medicine, Lund University, Lund, Sweden; Department of Immunotechnology, Science for Life laboratory, Lund University, Lund, Sweden

## Abstract

BRCA-deficient high-grade serous ovarian cancer is characterized by profound genomic instability and elevated replication-associated DNA damage, rendering these tumors initially sensitive to platinum-based chemotherapy and PARP inhibition. However, despite this vulnerability, most patients ultimately develop resistance, underscoring the need for therapeutic strategies that extend beyond DNA repair–targeted mechanisms.

Here, we introduce the MTDH–SND1 complex as a complementary therapeutic target that may expose additional stress vulnerabilities in ovarian cancer cells. We show that pharmacological disruption of the MTDH–SND1 interaction using C26-A6 increases susceptibility to ferroptosis-associated stress, an iron-dependent form of regulated cell death and that BRCA-deficient models are particularly more sensitive to this perturbation.

Notably, when combined with PARP inhibition, MTDH–SND1 disruption is associated with increased MHC class I expression in tumor cells, suggesting enhanced tumor visibility to the immune system. Together, these findings support a combination strategy that couples DNA repair disruption with metabolic and immunogenic remodeling in BRCA-deficient ovarian cancer.

## Introduction

High-grade serous ovarian carcinoma (HGSOC), the most common histologic subtype of epithelial ovarian cancer (EOC), accounts for approximately 70–80% of EOC-related deaths and remains the most lethal gynecologic malignancy. Standard treatment consists of cytoreductive surgery followed by platinum–taxane chemotherapy^1,2^, however, despite high initial response rates, the majority of patients experience recurrence and poor long-term outcomes^2^. Genetic predisposition plays a key role in EOC, particularly in HGSOC subtypes, with germline BRCA1 and BRCA2 mutations representing some of the most significant inherited risk factors^3,4^. Loss of BRCA1/2 function compromises homologous recombination (HR)–mediated DNA repair, creating a therapeutic vulnerability that is effectively targeted by poly (ADP-ribose) polymerase inhibitors (PARPi). Consequently, PARPi maintenance therapy has become standard of care for BRCA-mutant tumors following platinum response. However, many patients derive only limited benefit from PARPi monotherapy, and both intrinsic and acquired resistance frequently occur, even in tumors harboring BRCA mutations^5,6^. These clinical limitations highlight the need to expand therapeutic strategies in HGSOC by identifying new molecular targets and developing more effective targeted or combination treatments^7^.

Therapeutic responses in HGSOC are strongly influenced by BRCA1/2 mutation status, suggesting that BRCA1/2 loss may create context-specific vulnerabilities beyond defective DNA repair^8^. Recent studies suggest that BRCA1-deficient tumors exhibit altered sensitivity to ferroptosis, with changes in lipid peroxide detoxification pathways rendering them more susceptible to ferroptosis inducers, particularly in combination with PARP inhibition^9,10^. Ferroptosis has emerged as a promising therapeutic vulnerability in cancer, particularly in tumors that are resistant to apoptosis and conventional therapies. This form of regulated cell death is driven by the accumulation of iron-dependent lipid peroxides and is distinct from apoptosis, necrosis, and autophagy^11,12^.

The intrinsic ferroptosis vulnerability of BRCA1-deficient tumors creates a compelling rationale for combining ferroptosis-modulating agents with PARPi to achieve greater therapeutic benefit^9^. Leveraging this vulnerability alongside PARP inhibition may further enhance treatment efficacy. In this context, we previously demonstrated that targeting the MTDH–SND1 complex, can increase the sensitivity of ovarian cancer cells to ferroptosis^13^.

The MTDH–SND1 has also been implicated in tumor progression and resistance to therapy^14^, suggesting that it may function at the intersection of survival signaling and stress response pathways. C26-A6 is a small molecule that directly targets the MTDH–SND1 complex and its downstream pathways, effectively disrupting MTDH-mediated mechanisms that drive metastasis and resistance to therapy. Preclinical studies have demonstrated its efficacy in reducing tumor progression both as a monotherapy and in combination with immunotherapy^15,16^. However, whether targeting the MTDH–SND1 complex can potentiate ferroptosis and enhance PARP inhibitor responses in BRCA-deficient HGSOC remains unknown.

To evaluate the therapeutic potential of C26-A6 in combination treatments, we employed a pair of recently developed, genetically defined murine HGSOC cell line models. Designed to mimic homologous recombination-deficient/proficient tumors, BPPNM carries key mutations in TP53, BRCA1, PTEN, and NF1, along with MYC overexpression and PPNM shares the same alterations but retains wild-type BRCA1, serving as the HR-proficient counterpart^17^. These models enable studies in immunocompetent settings, allowing investigations into tumor microenvironment interactions, genetic alterations, and drug sensitivity. Leveraging these complementary models, this study explores tumor biology and potential novel therapeutic strategies in HGSOC by examining whether BRCA1 deficiency confers enhanced sensitivity to C26-A6 and by evaluating the therapeutic potential of combining MTDH–SND1 complex disruption with ferroptosis-inducing agents.

Furthermore, to evaluate the therapeutic potential of combining C26-A6 with PARP inhibition *in vivo*, we established HGSOC tumors using the BPPNM model in immunocompetent mice and assessed responses to the PARP inhibitor olaparib and C26-A6 administered alone or in combination. This *in vivo* approach was complemented by cell-based assays and multi-omic analyses, including proteomics, phosphoproteomics, and single-cell RNA sequencing, to elucidate the molecular mechanisms underlying treatment responses. Notably, combined C26-A6 and PARP inhibition resulted in enhanced immune activation within cancer cells, suggesting a possible interplay between MTDH–SND1 disruption, ferroptotic signaling, and BRCA1 deficiency.

## Results

### Combination effects of C26-A6 and ferroptosis inducers in ovarian cancer models

To evaluate the effect of C26-A6 on susceptibility of ovarian cancer cells to ferroptosis, we treated murine high grade ovarian tumor cell lines (BPPNM, PPNM) with C26-A6, either alone or in combination with the ferroptosis inducers erastin a system X_c_^-^ inhibitor^18^ or ML-162 a GPX4 inhibitor^19^ (Fig.1). Treatments were applied individually or in combination, and their effects on cell viability and clonogenic growth were assessed using CellTiter-Glo and colony formation assays.

**Figure 1.**
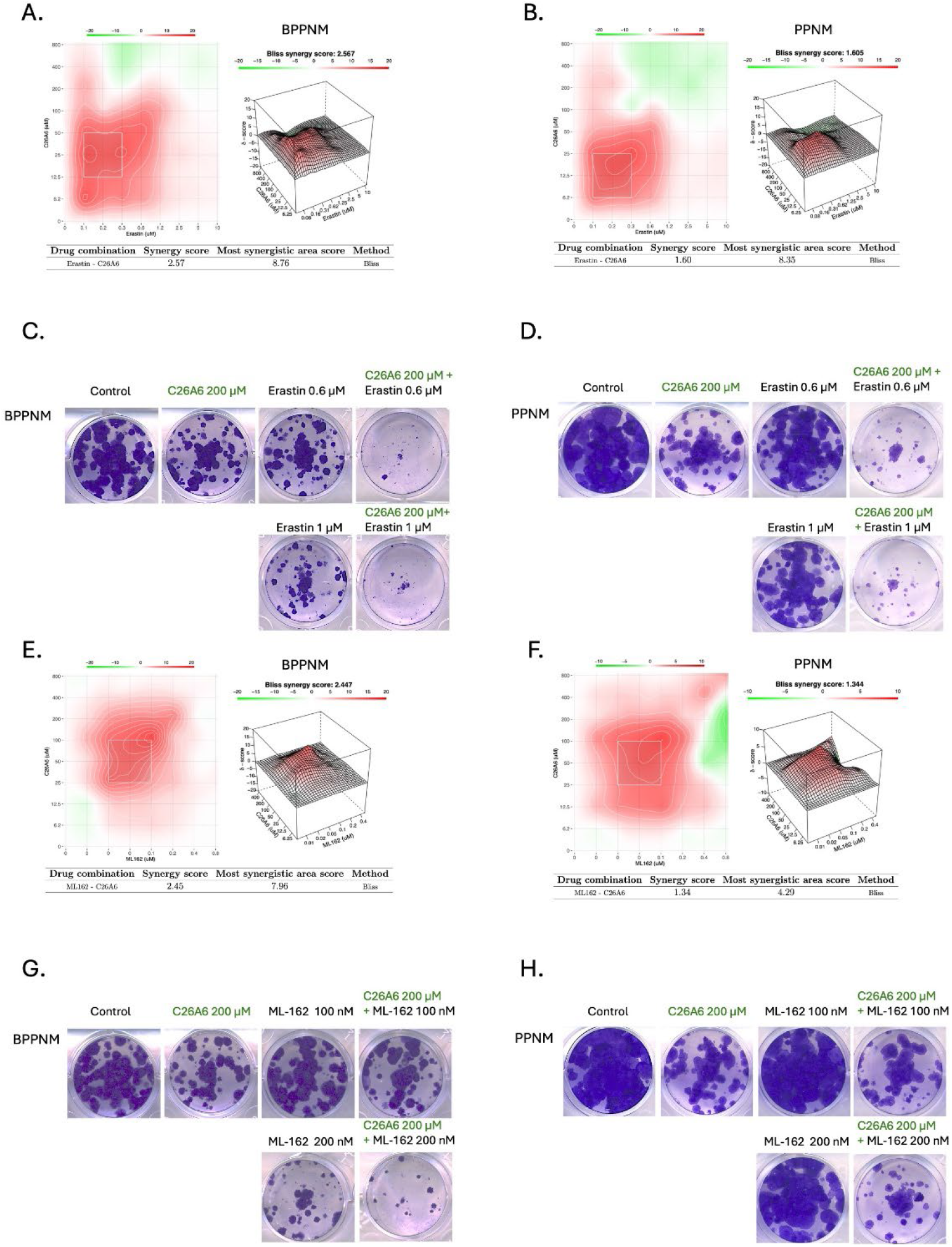
Disruption of the MTDH–SND1 axis enhances sensitivity of ovarian cancer cells to ferroptosis inducers. **A, B**. Bliss synergy analysis of C26-A6 in combination with erastin in BPPNM **A**. and PPNM **B**. cells. **C, D**. Colony formation assays following treatment with C26-A6 (200 μM), erastin (0.6 or 1 μM), or their combination in BPPNM **C**. and PPNM **D**. cells. **E, F**. Bliss synergy analysis of C26-A6 in combination with ML-162 in BPPNM **E**. and PPNM **F**. cells. **G, H**. Colony formation assays following treatment with C26-A6 (200 μM), ML-162 (100 or 200 nM), or the combination in BPPNM **G**. and PPNM **H**. cells.

Drug interaction analysis using the Bliss independence model^20^ revealed positive synergistic interactions between C26-A6 and erastin in both BPPNM and PPNM cells. The Most Synergistic Area Score (MSAS) was 8.76 in BPPNM and 8.35 in PPNM, with an overall synergy score of 2.57 and 1.60, respectively (Fig. 1A, B). Consistent with these findings, colony formation was markedly reduced by the combination of C26-A6 and erastin compared with either agent alone in both cell lines, with erastin exerting a stronger inhibitory effect in BPPNM than in PPNM cells (Fig. 1C, D). A similar interaction pattern was observed for the combination of C26-A6 with ML-162 in both cell lines (Fig. 1E, F). Correspondingly, combined treatment with C26-A6 and ML-162 significantly suppressed colony formation relative to either single-agent treatment in both cell lines, again with greater sensitivity observed in BPPNM cells (Fig. 1G, H). Together, these results suggesting that disruption of the MTDH–SND1 axis increases susceptibility to ferroptosis-inducing compounds, with the largest effects in BRCA1-deficient cells.

To investigate whether these combination effects extend to human ovarian cancer models, we repeated the treatments on the HGSOC cell line PEO1. Treatment with C26-A6, erastin or Ml-162 and their combinations resulted in MSAS of 9.13 and 15.63 in combination with erastin or Ml-162 respectively (Supplementary Fig. 1A, C), and significant decreases in colony formation under combined treatment conditions (Supplementary Fig. 1B, D). Synergy analysis in PEO1 cells indicated higher synergy scores, especially with ML-162 (overall synergy score: 4.53) compared to erastin (overall synergy score: 2.91), further supporting enhanced ferroptosis sensitivity in HRD cells.

While we previously evaluated C26-A6 sensitivity and its synergistic effects with ferroptosis inducers in additional high-grade serous ovarian cancer cell lines—including KURAMOCHI, CAOV3, and IGROV1^13^ we here used PEO1 due to its well-characterized BRCA2 deficiency. The inclusion of this BRCA-mutant model strengthens the relevance of our findings to homologous recombination-deficient (HRD) tumors. These results further support the hypothesis that HR deficiency enhances ferroptosis sensitivity and suggest that dual targeting of the MTDH–SND1 axis and ferroptosis pathways may represent a promising therapeutic strategy in BRCA-mutant ovarian cancer.

### Molecular mechanisms underlying the combination treatment in mice model

To investigate the molecular mechanisms underlying the combination effects of C26-A6 and ferroptosis inducers, we performed full proteomic profiling in two genetically defined murine HGSOC models: BPPNM and PPNM. These models enabled a controlled comparison of BRCA1-dependent signaling responses to treatment. Two cell lines were treated with C26-A6, erastin, or their combination, and samples were subjected to quantitative mass spectrometry– based proteomic analyses.

In a preliminary proteomic comparison between the BPPNM and PPNM cell lines, 628 proteins were significantly different without treatment (FDR < 0.05 and |log2 fold change| > 1). Among those 304 were higher abundance in BPPNM and 324 were higher in PPNM. Over-representation analysis revealed that PPNM cells exhibited increased expression of proteins involved in cell adhesion, junctional architecture, and homeostasis. In contrast, BPPNM cells showed elevated activity in pathways related to fatty acid metabolism, metabolic, and lysosomal function (Fig. 2A).

**Figure 2.**
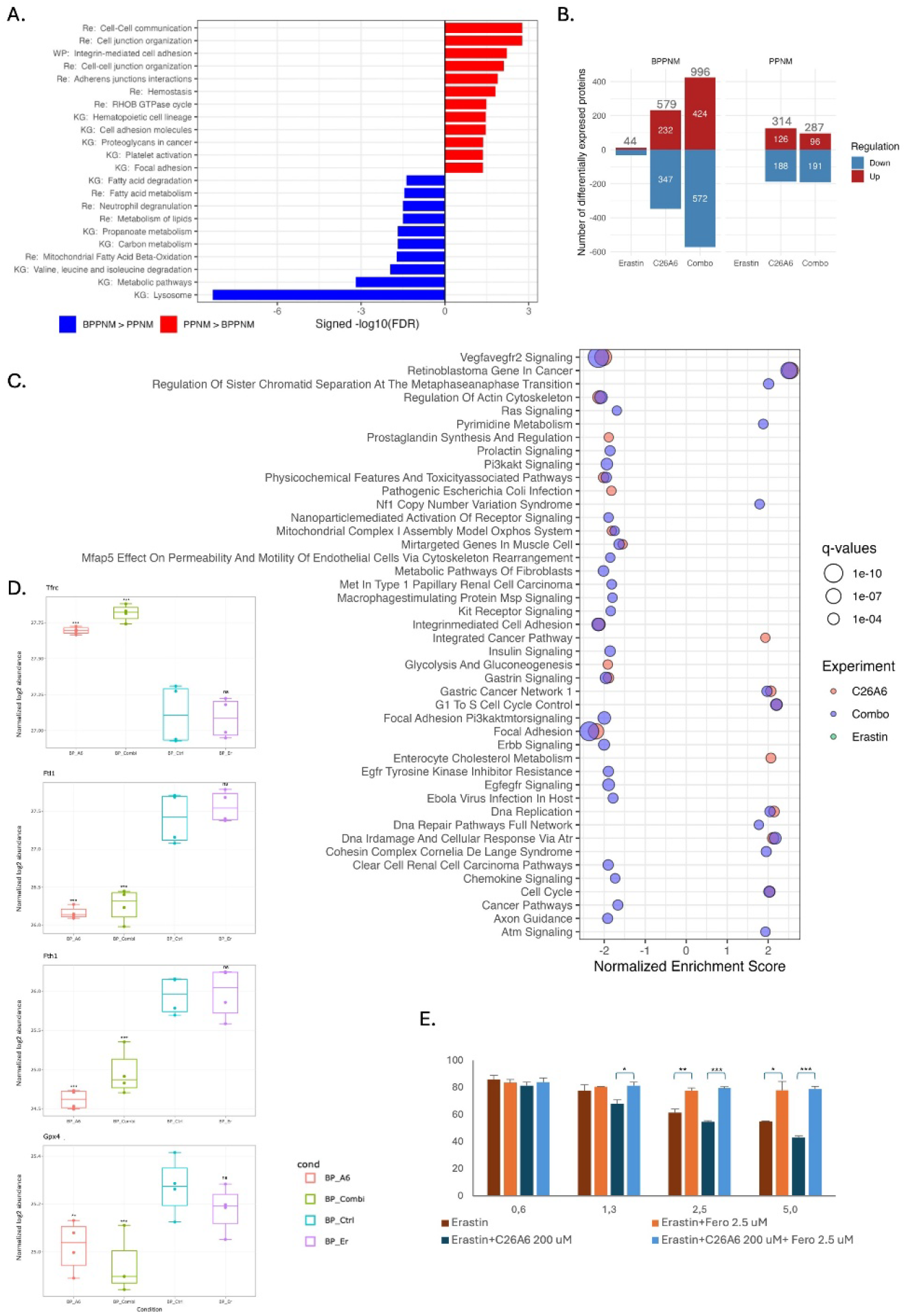
Proteomic profiling reveals differential pathway enrichment in BPPNM and PPNM. **A**. Diverging bar plot of KEGG (KG), Reactome (Re), and WikiPathways (WP) enrichment comparing PPNM and BPPNM. Bars to the right (red) indicate pathways enriched in PPNM, bars to the left (blue) indicate pathways enriched in BPPNM, with bar length representing −log10(false discovery rate). **B**. Bar graphs displaying the number of DEPs (FDR < 0.05) in BPPNM and PPNM under different treatment conditions compared to untreated controls. **C**. GSEA based on WikiPathways of BPPNM across all treatment conditions, with pathways shown at a significance threshold of pvalueCutoff < 0.01. **D**. Box plots showing the abundance of TFRC, FTL1, FTH1, and GPX4 protein expression in BPPNM cells under erastin, C26-A6, combination, and untreated conditions. **E**. Rescue assay with ferrostatin-1. Cell viability values were normalized to the corresponding untreated or ferrostatin-1–only control groups and expressed as percentage (%). Experiments were performed in triplicate (n = 3); error bars represent the standard error of the mean (SEM). Statistical significance was determined using an unpaired two-tailed Student’s t-test. Significance is indicated as follows: ns, not significant; *p* < 0.05 (*), *p* < 0.01 (**), *p* < 0.001 (***). All comparisons are relative to the control group.

To assess early drug responses while avoiding overt cytotoxicity, we applied low doses (IC20) of each compound, aiming to detect initial signs of sensitivity without inducing full cell death. Comparison of treatment effects using C26-A6, erastin, and their combination revealed greater efficacy in BPPNM cells compared to PPNM cells (Fig. 2B). In BPPNM cells, treatment with C26-A6 resulted in 579 significantly differentially expressed proteins (DEPs; FDR < 0.05), while erastin alone led to only 44 DEPs. Remarkably, the combination treatment yielded 996 DEPs, suggesting a greater-than-additive effect when both drugs are used together. In contrast, PPNM cells showed no significant DEPs in response to erastin alone. Treatment with C26-A6 and the combination resulted in 314 and 287 DEPs, respectively, indicating no clear combination effect at these low doses in the BRCA-proficient context. Thus, BRCA1 deficiency enhances proteomic responsiveness to combined ferroptosis- and MTDH-SND1 interaction-targeting treatments, supporting a potential combination effect between ferroptosis induction and DNA repair deficiency.

Gene set enrichment analysis (GSEA)^21^ was performed using WikiPathways gene sets on BPPNM proteomics data, visualized via a bubble plot, to compare pathway enrichment across treatment conditions (Fig. 2C). While erastin alone did not yield pathway enrichment at the applied cutoff, the combination treatment produced a broader enrichment profile than C26-A6 alone. This pattern is consistent with the combination engaging pathways that were not detected with either agent alone at these doses.

Within these enriched pathways, the most significantly regulated pathways shared between C26-A6, and combination treatment included VEGFA signaling and focal adhesion pathways, as well as retinoblastoma-associated gene set, with both treatments showing regulation in the same direction. However, the combination treatment uniquely suppressed a broader set of pro-survival and proliferative signaling pathways, including RAS, PI3K–AKT, insulin, EGFR, and ErbB signaling, as well as key regulators of cellular metabolism such as fibroblast metabolic programs, MET signaling, and nanoparticle-mediated receptor activity.

This widespread repression of growth-supportive signaling suggests that the combination treatment is associated with increased molecular stress and increased susceptibility to cell death–associated processes, including ferroptosis (Fig. 2D), in a genetically vulnerable context like BRCA1 deficiency. Disruption of MTDH-SND1 interaction in BPPNM cells was associated with lower abundance of the ferritin subunits FTH1, FTL1, and GPX4, together with higher abundance of the transferrin receptor TFRC. This pattern is compatible with reduced iron sequestration and reduced capacity for lipid peroxide detoxification, hallmark features of ferroptosis^22^. Notably, a similar marker pattern was observed in the BRCA-proficient cell line (Supplementary Fig. 2A), suggesting that this response may occur outside BRCA1 deficiency. In parallel, pathway enrichment analysis suggested enrichment of NRF2-related pathways in PPNM cells (Supplementary Fig. 2B), consistent with an antioxidant stress exposure^23^. Importantly, ferroptotic cell death was functionally validated using a rescue assay (Fig. 2E), in which treatment with the ferroptosis inhibitor ferrostatin-1 fully rescued cell death induced by erastin alone as well as by the erastin and C26-A6 combination. The comparable rescue efficiency indicates that the combination-induced cytotoxicity remains ferroptosis-dependent. Collectively, these results indicate that the C26-A6 sensitizes cancer cells to ferroptotic cell death.

### Molecular mechanisms underlying the combination treatment in human model

Comparison of treatment effects using C26-A6, erastin, and their combination in human high-grade serous ovarian cancer cell line PEO1 is shown in Fig. 3. The combination treatment yielded a substantially greater number of DEPs, 1913 (FDR < 0.05), compared with 1156 for C26-A6 alone and 24 for erastin alone. These results mirror the findings in mouse BPPNM cells, further supporting the enhanced molecular impact of the combination treatment, particularly in BRCA-deficient contexts such as PEO1.

**Figure 3.**
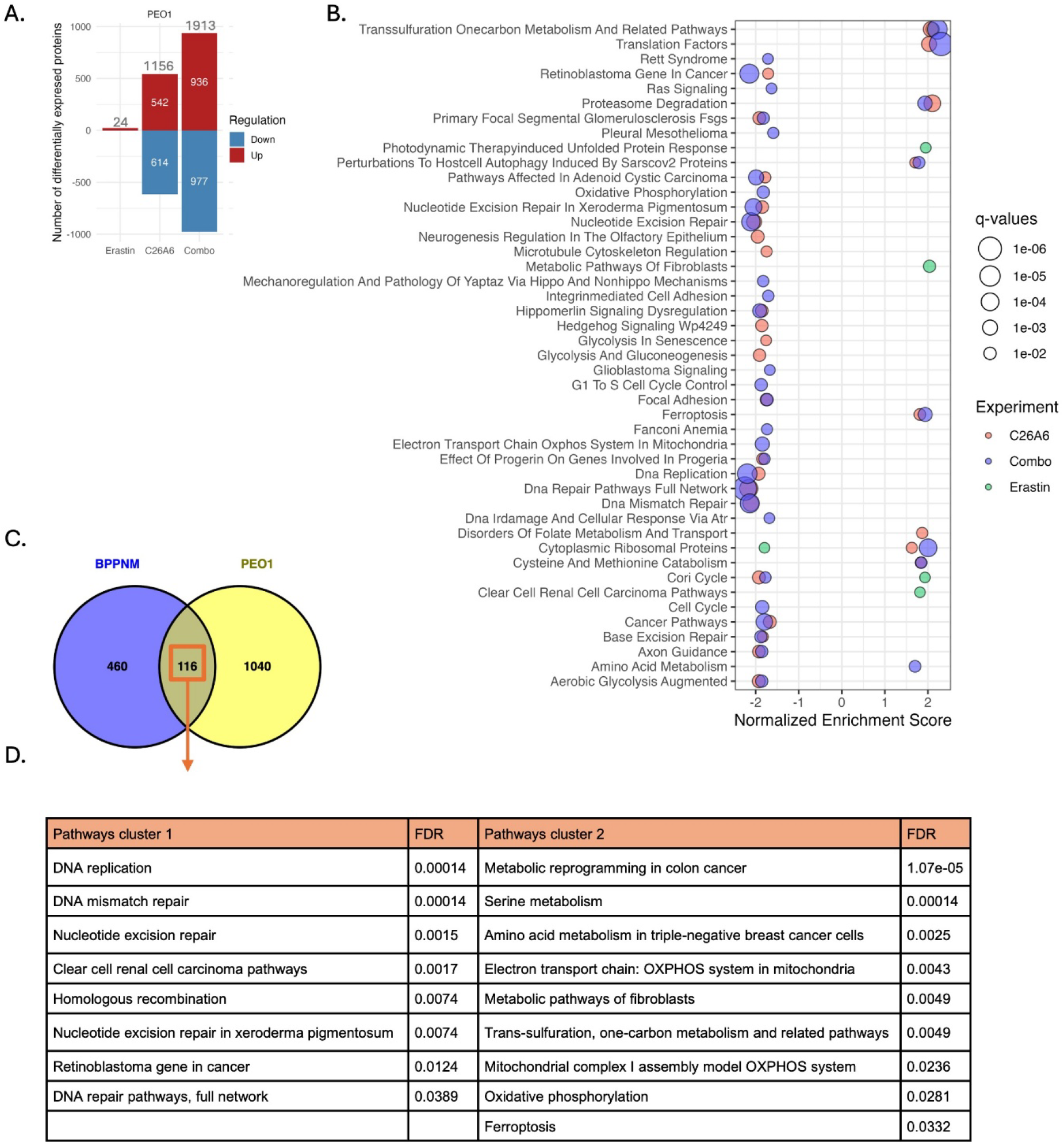
Proteomic profiling reveals differential pathway enrichment in PEO1 cell line. **A**. Bar graph showing the number of differentially expressed proteins (DEPs; FDR < 0.05) in PEO1 cells following treatment with erastin, C26-A6 or their combination, compared to untreated controls. **B**. GSEA based on WikiPathways of PEO1 across all treatment conditions. Enriched pathways are shown at a significance threshold of *p* < 0.01. **C**. Venn diagram showing the overlap of DEPs (FDR < 0.05) across two cell lines (BPPNM and PEO1) under C26-A6 treatment. **D**. Functional enrichment analysis of the shared 116 DEPs in WikiPathways databases.

In PEO1 cells, combination treatment was associated with downregulation of multiple mitogenic, metabolic, and DNA repair–related pathways, including RAS, integrin signaling, oxidative phosphorylation (OXPHOS), and ATR signaling. Similarly, in both BPPNM and PEO1 cells, combination treatment resulted in dysregulation of major pro-survival, metabolic, and DNA damage response pathways. In both models, key proliferative and metabolic networks, including RAS signaling, cell adhesion pathways, and mitochondrial respiration, were suppressed, alongside disruption of DNA damage response pathways through ATR signaling. In addition, pathway enrichment analysis indicated activation of ferroptosis-associated pathways following combination treatment.

Next, to uncover the shared effects of C26-A6 on BRCA-deficient models, we first translated the mouse DEPs (FDR < 0.05) from BPPNM to their *Homo sapiens* orthologs^24^ (Supplementary Data.1). We then compared them with DEPs from PEO1 cell lines, identifying 116 shared DEPs (Fig. 3C, Supplementary Data. 2), which were subjected to targeted pathway enrichment analysis using STRING^25^ Pathway enrichment analysis revealed two major group of affected pathways (Fig. 3D). One group comprised alterations in DNA replication and DNA repair pathways, including homologous recombination, mismatch repair, and nucleotide excision repair. In parallel, a second group included several mitochondrial and metabolic pathways, notably oxidative phosphorylation, mitochondrial complex I assembly, serine and amino acid metabolism, one-carbon and trans-sulfuration metabolism. Importantly, ferroptosis-related pathways were also enriched. These findings suggest that MTDH-SND1 distruption induces a coordinated interference of genome maintenance and mitochondrial metabolism in BRCA-deficient cancer cells.

Thus far, we have demonstrated that C26-A6 exerts a dose-dependent cytotoxic effect across both murine and human ovarian cancer cell lines, irrespective of BRCA status. *In vitro* studies revealed that C26-A6 enhances susceptibility to ferroptosis, and its combination with the ferroptosis inducer erastin resulted in a clear combination response. Furthermore, pathway enrichment analysis of shared DEPs in BRCA-deficient models revealed not only mitochondrial dysfunction and metabolic rewiring, but also significant alterations in homologous recombination and DNA repair pathways. These findings raised the possibility that targeting MTDH-SND1 interaction may sensitize BRCA-deficient cells not only to ferroptosis, but also to PARP inhibition. To explore this possibility, we first evaluated the effects of combined C26-A6 and olaparib treatment *in vitro*. Drug interaction analysis revealed a good combination effect in BPPNM cells, with a MSAS of 6.93 (Supplementary Fig. 3A), whereas only minimal score was observed in the BRCA-proficient PPNM cells (MSAS 0.93, Supplementary Fig. 3B). Consistent with this, combined treatment resulted in a marked reduction in clonogenic survival in BPPNM cells compared with either agent alone (Supplementary Fig. 3C).

### Targeting MTDH-SND1 enhances the efficacy of PARP inhibition *in vivo*

To test this hypothesis *in vivo*, we established an immunocompetent ovarian cancer mouse model using the BRCA1-deficient BPPNM cell line. Mice were treated with C26-A6 alone, olaparib alone, or a combination of both, enabling us to evaluate whether C26-A6 could enhance the therapeutic efficacy of PARP inhibition in BRCA1-deficient tumors.

As shown in Fig. 4A, four groups of C57BL/6 mice were intraperitoneally injected with the BRCA1-deficient ovarian cancer cell line BPPNM. Three weeks post-injection, once tumors were palpable through the skin, treatment was initiated. Mice received C26-A6 (5 times per week)^15^, olaparib (2 times per week)^17^, the combination of both (at the same dosing schedules), or vehicle control. Treatments continued for one month. While neither C26-A6 nor olaparib alone significantly reduced tumor size or abdominal area compared to vehicle, the combination treatment resulted in a significant reduction in both tumor size and abdominal swelling (Fig. 4B-D, Supplementary Data. 4). These findings highlight the combination effect of C26-A6 and olaparib in suppressing tumor growth in BRCA1-deficient ovarian cancer *in vivo*.

**Figure 4.**
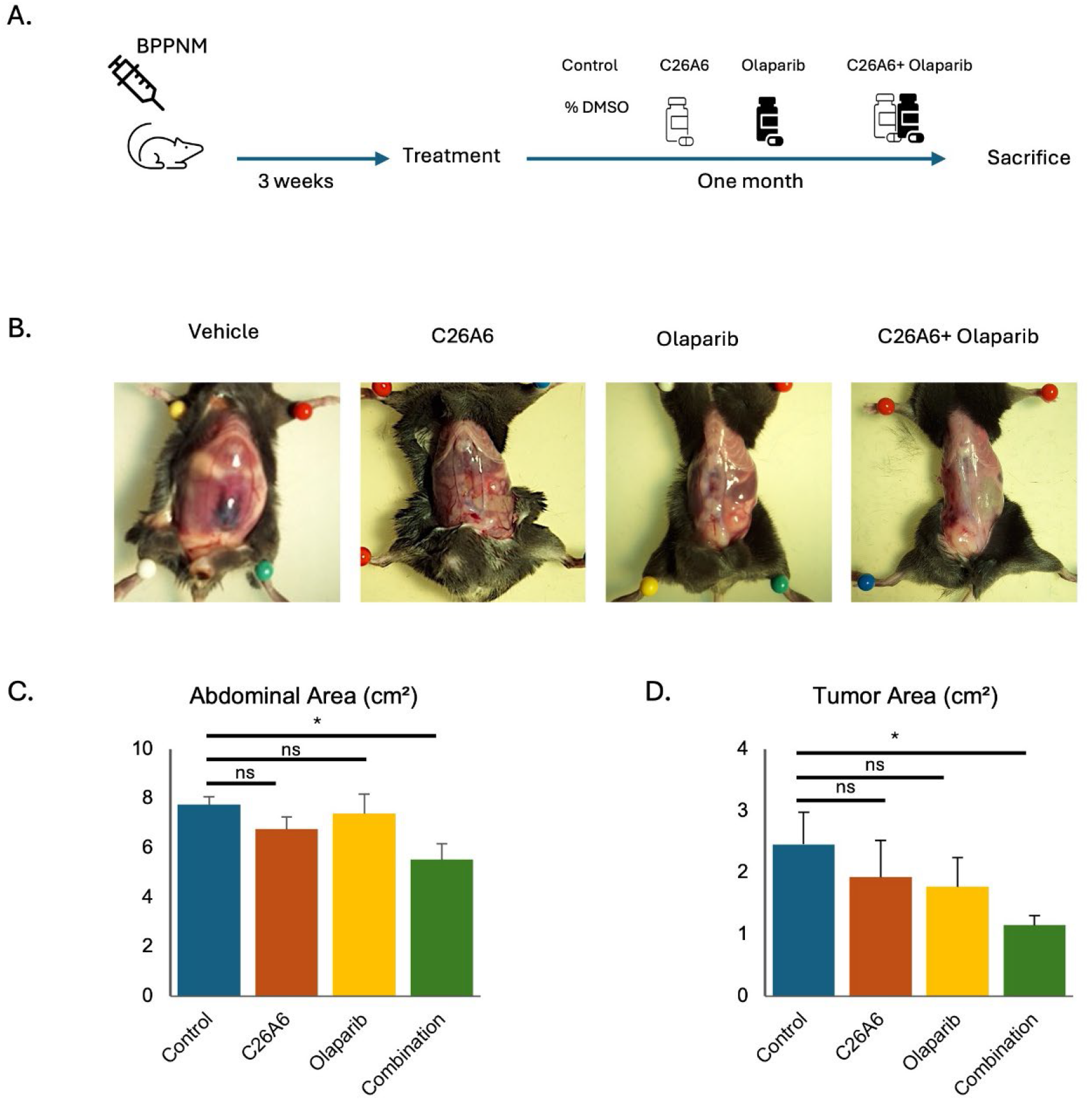
*In vivo* assessment of C26-A6 treatment in an immunocompetent mouse model. **A**. Schematic of the experimental treatment strategy. C57BL/6 mice were intraperitoneally injected with BPPNM cells and treated according to the indicated protocol. **B**. Representative images showing ascites accumulation and abdominal enlargement in mice following tumor development. **C**. Quantification of abdominal area and **D**. tumor burden across all experimental replicates. Bar graphs show mean values; error bars represent standard error of the mean (SEM). Statistical significance was determined using an unpaired two-tailed Welch’s t-test comparing treatments with controls. Significance levels are indicated as: ns, not significant; *p* <0.05 (*).

### Combination treatment induces distinct proteomic and phosphoproteomic signatures *in vivo*

To investigate the molecular changes underlying the enhanced therapeutic effect, we performed mass spectrometry-based bulk proteomics and phosphoproteomics on tumor tissues collected from the four treatment groups. Proteomics analysis revealed a higher number of DEPs (=812) in tumors from the combination treatment group compared to those treated with C26-A6 or olaparib alone (Fig. 5A). GSEA of the proteomic data revealed broader and more pronounced pathway enrichment in the combination-treated tumors relative to single-agent treatments (Fig. 5B). These findings indicate that combination therapy induces more extensive remodeling of protein expression and signaling networks *in vivo*, consistent with its enhanced antitumor activity.

**Figure 5.**
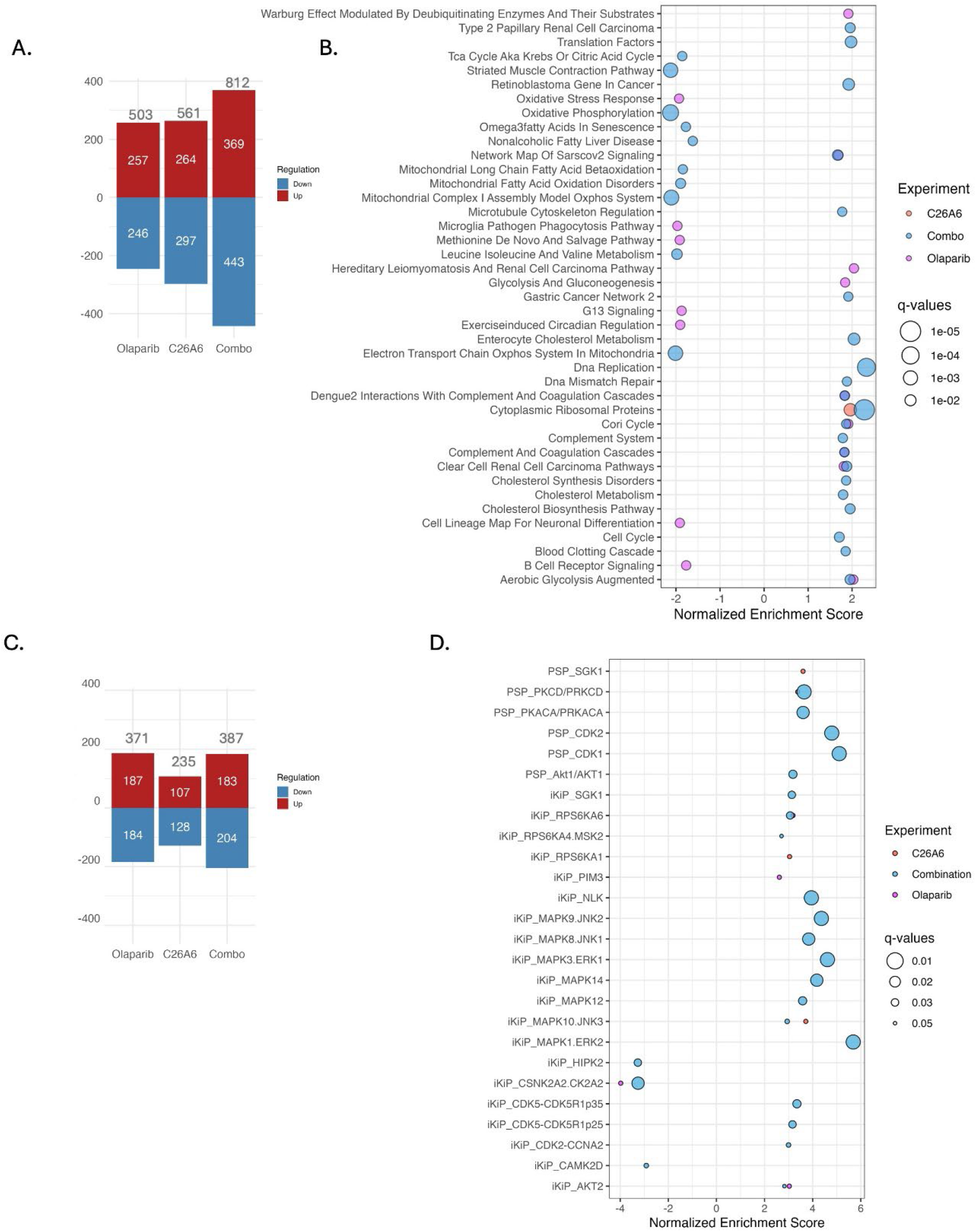
Proteomic and phosphoproteomic analysis of treatment-specific pathway alterations in tumor tissues. **A**. Number of DEPs (*p* < 0.05) identified in response to olaparib, C26-A6, and their combination compared to untreated control. **B**. GSEA based on WikiPathways showing pathway enrichment in response to C26-A6, olaparib, and combination treatment. Pathways are shown at a significance threshold of FDR < 0.05. **C**. Number of differentially expressed Phosphopeptides (DEPPs; *p* < 0.05) identified in response to olaparib, C26-A6, and their combination compared to untreated control. **D**. Kinase enrichment analysis using PTM-SEA based on phosphoproteomic data from tumor tissues treated with C26-A6, olaparib, or their combination. Significantly enriched kinases were identified using an FDR cutoff of < 0.05.

Specifically, GSEA of the proteomic data demonstrated significant negative enrichment of pathways associated with mitochondrial energy metabolism, including oxidative phosphorylation and fatty acid oxidation, in tumors treated with the combination of C26-A6 and olaparib. In contrast, pathways related to protein synthesis and cholesterol biosynthesis showed positive enrichment in the same treatment group. Additionally, complement-associated pathways were enriched, indicating altered inflammatory or tumor microenvironment–related signaling. Collectively, these results highlight extensive metabolic and signaling pathway reprogramming in response to combination treatment.

To further investigate the signaling responses induced by treatment, we performed mass spectrometry–based phosphoproteomic analysis on tumor tissues from the same treatment groups. In contrast to the proteomic analysis, phosphoproteomic profiling identified a more limited number of differentially expressed phosphopeptides (=387) in tumors from the combination treatment group compared to those treated with C26-A6 or olaparib alone (Fig. 5C). To assess upstream signaling alterations, we subsequently applied post-translational modification set enrichment analysis (PTM-SEA)^26^ to the phosphoproteomic dataset (Fig. 5D). Due to limitations in the mouse-specific PTM libraries, we used the human kinase substrate database for kinase activity inference. Interestingly, significant changes in kinase activity were observed only in the combination treatment group, whereas minimal enrichment was observed in tumors treated with single agents.

PTM-SEA of the phosphoproteomic dataset revealed increased inferred activity of multiple stress- and cell cycle–associated kinases in tumors treated with the combination of C26-A6 and olaparib, including JNK1/2 (MAPK8/9), ERK1/2 (MAPK3/1), p38 (MAPK14), CDK1/2/5, AKT1, and SGK1/3 (Fig. 5D). In contrast, reduced inferred activity was observed for HIPK2, a kinase implicated in DNA damage response and apoptosis, as well as CK2A2, which has been linked to proliferative and survival signaling. Together, these phosphorylation changes indicate broad alterations in stress- and cell cycle–related signaling pathways in response to combination treatment. As these analyses were performed on bulk tumor tissue, the observed signaling changes reflect an aggregate response of the tumor microenvironment and malignant cells rather than cell type–specific effects.

### ScRNA-seq reveals increased epithelial antigen presentation under combination therapy

To examine cell type–specific responses associated with the bulk proteomic and phosphoproteomic alterations observed *in vivo*, we performed single-cell RNA sequencing (scRNA-seq) on tumor tissues from control, C26-A6–treated, olaparib-treated, and combination-treated mice. Tumor samples from one mouse per treatment group were processed for scRNA-seq analysis. Following quality control and integration, unsupervised clustering identified transcriptionally distinct cell populations within the tumor microenvironment, which were visualized by uniform manifold approximation and projection (UMAP) (Fig. 6A). Cell identities were assigned using canonical markers and cross-validated with Celltypist and GSEA. This approach identified eight major cellular compartments, including epithelial/cancer cells, stromal populations comprising cancer-associated fibroblasts (CAFs) and ECM-remodeling CAFs, endothelial cells, pericytes, myeloid cells, proliferating cells, and T/NK cells. B cells were excluded from downstream analyses due to low abundance. Cell-type distributions showed some minor shifts across samples, suggesting that cell-type abundance could be altered between the treatment groups at this time point (Supplementary Table 1). However, proteomics-based cell-type distribution estimates through data deconvolution across all replicates indicated that these changes were within natural tumor heterogeneity and not treatment-specific (Supplementary Fig. 4).

**Figure 6.**
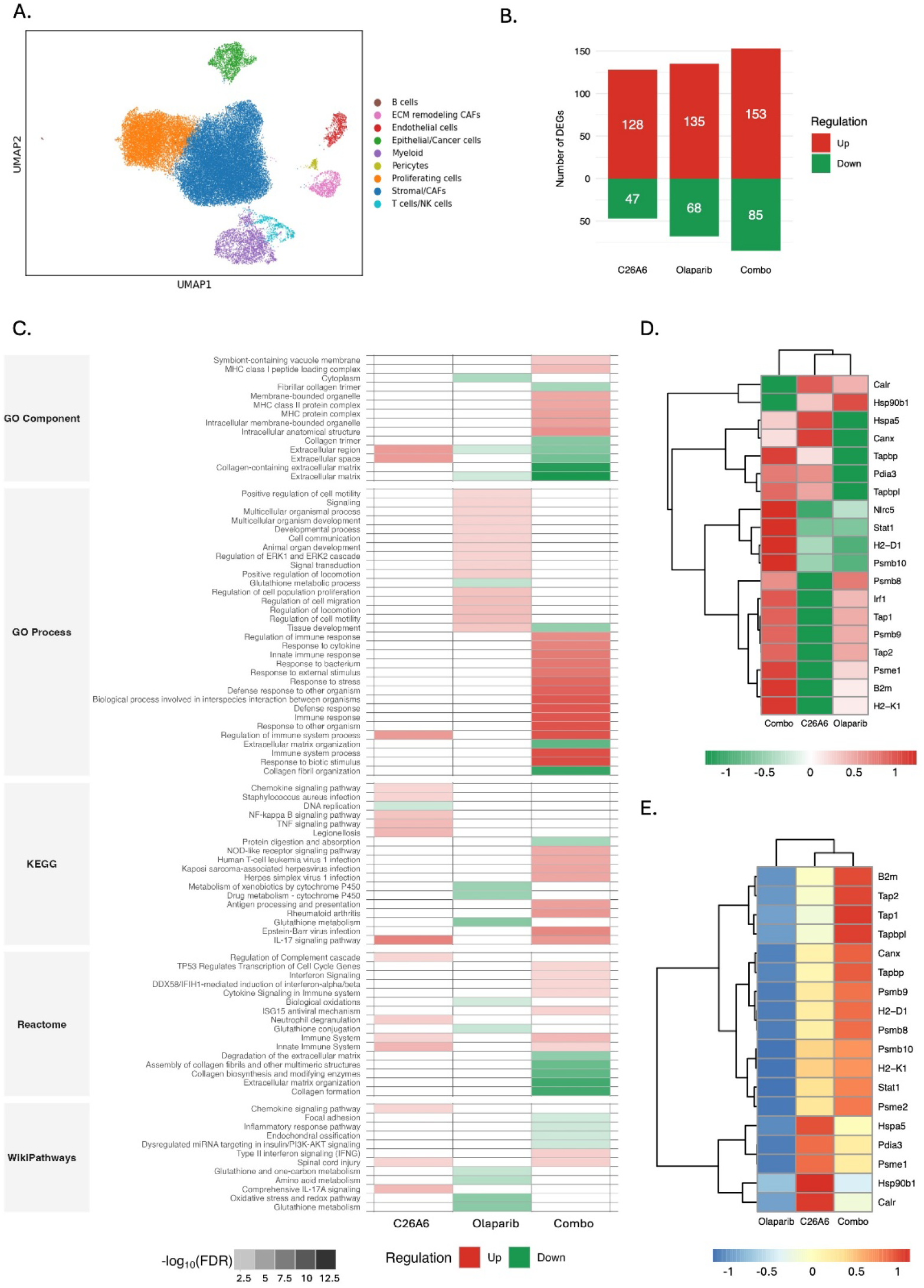
Single-cell RNA sequencing analysis. **A**. UMAP visualization of scRNA-seq cell clusters from tumor tissue **B**. Differentially expressed genes in epithelial tumor cells across treatment groups (FDR < 0.05, |log_2_FC| ≥ 1). **C**. Over-representation analysis of epithelial tumor cell differentially expressed genes Pathways are shown at a significance threshold of FDR < 0.05. **D**. Heatmap of MHC class I antigen processing and presentation–related gene expression in epithelial tumor cells across treatment groups (|log_2_FC| ≥ 1). **E**. Heatmap of MHC class I–associated protein expression from bulk proteomic analysis across treatment groups (|log_2_FC| ≥ 1). In proteomic analyses, red and blue indicate up- and downregulated proteins, respectively, whereas in scRNA-seq analyses, red and green indicate up- and downregulated genes.

Although malignant epithelial cells constituted a smaller proportion of the total cell population compared with stromal and proliferating compartments, downstream analyses were focused on the epithelial cancer cell cluster. This decision was motivated by the central role of malignant cells in mediating treatment response and by the objective of identifying tumor cell–intrinsic transcriptional changes associated with combination therapy. Accordingly, differential gene expression analysis was performed within the epithelial compartment. The combination treatment group exhibited a higher number of differentially expressed genes (DEGs) compared with single-agent treatments (FDR < 0.05, |log_2_FC| ≥ 1, Fig. 6B). To functionally characterize these transcriptional changes, over-representation analysis was conducted using multiple pathway databases (Fig. 6C, Supplementary Data. 3). This analysis revealed broader pathway enrichment in the combination treatment group, with notable upregulation of immune-related pathways, including antigen processing and presentation, immune system processes, and MHC protein complex–associated pathways. In contrast, the most prominently downregulated pathways in this condition were related to collagen organization and extracellular matrix– associated processes. To further examine immune-related transcriptional changes within the epithelial tumor cell compartment, we next focused on genes involved in MHC class I– mediated antigen processing and presentation. Expression of MHC-I–associated genes were assessed across all treatment conditions (Fig. 6D), revealing significant upregulation of multiple antigen presentation–related genes in epithelial cells from combination-treated tumors compared with single-agent and control groups.

To determine whether these transcriptional changes were reflected at the protein level, we revisited the bulk proteomic dataset. Consistent with the scRNA-seq findings, protein expression of MHC-I–associated components were most prominently increased in tumors from the combination treatment group, with more modest upregulation observed following C26-A6 treatment and no detectable increase in olaparib-treated tumors (Fig. 6E). These findings suggest that combination therapy may enhance tumor cell–intrinsic antigen presentation capacity, potentially influencing tumor–immune interactions within the tumor microenvironment.

## Discussion

HGSOC is most often diagnosed at an advanced stage, when widespread dissemination and therapy resistance severely limit long-term clinical benefit from standard treatments such as surgery and platinum-based chemotherapy^1^. This underscores the need for therapeutic strategies that target tumor cell vulnerabilities beyond canonical DNA-damaging approaches. BRCA-deficient tumors, in particular, exhibit distinct metabolic and stress-response features that may be exploited through rational combination therapies. Recent studies have demonstrated that the interaction between MTDH and SND1 plays a crucial role in metastasis and drug resistance in various cancers^27,28^. In our previous work, we demonstrated that either treatment with C26-A6 or silencing of MTDH or SND1 individually led to dysregulation of ferroptosis-related proteins. Interestingly, combining C26-A6 with ferroptosis inducers enhanced antitumor effects^13^.

BRCA1 deficiency is associated with profound metabolic reprogramming that appears to prime tumor cells for ferroptosis-associated vulnerability^9,10^. Comparative proteomic profiling revealed enrichment of fatty acid metabolism, lipid handling, and broader metabolic pathways in BRCA1-deficient ovarian cancer cells, suggesting increased lipid turnover and remodeling relative to BRCA-proficient counterparts (Fig. 2A). Increased lipid metabolism—especially involving polyunsaturated fatty acids (PUFAs) and impaired or overloaded β-oxidation—can lead to a buildup of lipid peroxides, which are key drivers of ferroptosis^29,30^. In this context, the lysosome pathway emerged as the most strongly activated pathway in the BRCA-deficient model, together with increased mitochondrial activity and enhanced fatty acid and amino acid degradation. This coordinated proteomic profile is consistent with a ferroptosis-associated metabolic state in which lysosomes act as central hubs for iron handling, lipid turnover, and membrane damage responses—processes that have been linked to elevated lysosomal iron pools and increased vulnerability to iron-dependent lipid peroxidation^31^.

Consistent with the metabolic vulnerabilities observed in BRCA-deficient cells, ferroptosis inducers exerted greater antitumor activity in the BRCA-deficient model, supporting the notion that loss of BRCA function sensitizes tumor cells to ferroptotic stress^9^. Moreover, combining ferroptosis inducers with disruption of the MTDH–SND1 interaction further enhanced therapeutic efficacy in BRCA-deficient cells relative to BRCA-proficient counterparts, indicating that BRCA status modulates susceptibility to ferroptosis-driven cytotoxicity. Notably, the observation that ferrostatin-1 rescued cell death induced by the erastin and C26-A6 combination to a similar extent to erastin alone supports a ferroptosis-dependent mechanism and suggests that C26-A6 primarily enhances cellular susceptibility to ferroptotic stress

Across two independent BRCA-deficient models (BPPNM, PEO1), combination treatment induced a shared transcriptional response, despite differences in species origin. Although individual pathways exhibited cell line–specific directionality, the substantial overlap in enriched programs indicates that the combination therapy predominantly engages common stress-response mechanisms intrinsic to BRCA-deficient cells. In particular, combination treatment dysregulated multiple genes associated with DNA replication, cell cycle progression, and DNA damage response signaling, including ATR- and retinoblastoma (RB)–associated pathways, consistent with altered regulation of replication stress and cell cycle checkpoint processes. ATR signaling is especially important in homologous recombination–deficient cells, where it plays a key role in coordinating cellular responses to replication-associated stress^32^. Dysregulation of ATR- and RB-linked pathways therefore suggests that combination therapy amplifies replication-associated stress while simultaneously perturbing cell cycle control mechanisms, reinforcing a shared vulnerability across BRCA-deficient cell models.

In BRCA-deficient cell lines, C26-A6 appears to engage multiple, interconnected stress-response programs. Pathway-level analyses indicate that C26-A6 preferentially impacts two functional axes: genome maintenance and ferroptosis-associated metabolism. Encompassed DNA replication and DNA repair processes, consistent with the central role of BRCA in maintaining genomic stability^33^ and suggesting that C26-A6 may further perturbs replication-associated stress in this genetic background. In parallel, enrichment of ferroptosis-related and metabolic pathways, indicating that C26-A6 may also modulates redox and metabolic programs^29^ linked to lipid peroxidation and cellular stress responses. Together, these findings suggest that C26-A6 exerts dual effects in BRCA-deficient cells by simultaneously targeting genome maintenance pathways and ferroptosis-associated metabolic vulnerabilities.

Taken together, our findings indicate that BRCA-deficient tumors display coordinated dysregulation of DNA replication and repair pathways alongside ferroptosis-associated metabolic programs, which are further modulated by C26-A6 treatment. Given the established clinical sensitivity of BRCA-deficient cancers to PARP inhibition, these results provided a rationale to explore combination strategies that simultaneously target DNA repair and stress-adaptive pathways. However, resistance to PARP inhibitors can arise through multiple mechanisms, including restoration of homologous recombination via BRCA1/2 reversion mutations or loss of 53BP1, stabilization of stalled replication forks, and reduced PARP trapping^34,35^. These resistance pathways highlight the need for combination approaches and support the evaluation of PARP inhibitors together with agents targeting complementary vulnerabilities, such as immunotherapy^34-36^, or ferroptosis inducers^37^. Based on this, we next assessed the therapeutic efficacy of combining the PARP inhibitor olaparib with C26-A6 in immunocompetent mice to evaluate treatment responses in the context of an intact tumor microenvironment.

To model clinically relevant disease complexity, tumors were allowed to establish and disseminate prior to treatment initiation, approximating the advanced and metastatic stage at which HGSOC is typically diagnosed. Although olaparib is commonly administered as a maintenance therapy in the clinic following platinum-based chemotherapy^38^, our study aimed to explore whether combining olaparib with the ferroptosis-sensitizing compound C26-A6 could enhance therapeutic effects in a platinum-independent context. Our objective was to assess the initial therapeutic impact of the olaparib and C26-A6 combination on tumor burden and associated molecular signatures under aggressive disease conditions.

In the *in vivo* setting, combination therapy was associated with a pronounced reduction in tumor burden, reflected by decreased tumor size and abdominal involvement, indicating enhanced therapeutic efficacy compared to single-agent treatments. Consistent with these phenotypic effects, integrated proteomic and phosphoproteomic analyses revealed substantially broader molecular remodeling in combination-treated tumors. At the proteomic level, combination therapy suppressed mitochondrial metabolism and fatty acid oxidation while enriching pathways related to cholesterol biosynthesis and complement and coagulation cascades, collectively defining a metabolically stressed tumor state.

This coordinated metabolic profile is consistent with a metabolically stressed tumor state and reflects adaptive remodeling of energetic and biosynthetic pathways in response to energy limitation and cytotoxic therapeutic pressure^39^. Upregulation of complement-associated pathways in bulk tumor proteomics is consistent with treatment-induced tumor damage and activation of innate immune sensing mechanisms. Cytotoxic therapies such as radiotherapy and chemotherapy have been shown to trigger local complement activation as a consequence of tumor cell injury and membrane perturbation, thereby linking tissue damage to immune engagement^40,41^. Acute, localized complement activation can promote immunogenicity and support tumor clearance, whereas chronic or dysregulated signaling may instead contribute to immunosuppressive remodeling of the tumor microenvironment^41^. Thus, the selective enrichment of complement pathways observed only in the combination treatment group is consistent with a heightened therapy-induced stress and injury response characteristic of effective cytotoxic treatment.

Bulk phosphoproteomic analysis further revealed a distinct treatment-induced signaling state that was selectively observed in the combination therapy group but not in either single-agent condition. This state was characterized by predominant activation of MAP kinase pathways, including ERK and JNK, which are well-established mediators of redox stress sensing, cell cycle regulation, and adaptive survival signaling^42^. In contrast, kinase activity analysis indicated reduced activity of HIPK2, a kinase involved in apoptosis and DNA damage response^43^, and CK2A2, which supports proliferative and survival of cancer cells^44^. Together, these phosphoproteomic changes align with the proteomic evidence of metabolic stress and adaptive remodeling, indicating coordinated rewiring of metabolic and signaling networks specifically under combination treatment. As bulk proteomic and phosphoproteomic approaches cannot resolve the cellular origin of inflammatory and immune-associated signals, these findings likely reflect integrated responses from both malignant cells and the surrounding tumor microenvironment, highlighting the complexity of therapy-induced signaling adaptations *in vivo*.

To detect tumor cell–intrinsic responses from microenvironmental contributions, we applied single-cell RNA sequencing to explore transcriptional programs associated with combination therapy. The emergence of antigen presentation–related gene programs within cancer cells suggests that, in addition to inducing metabolic and signaling stress, combination treatment may alter the intrinsic immunogenic state of tumor cells. In line with this notion, we observed consistent upregulation of MHC class I–associated genes and proteins following combination therapy, indicating enhanced tumor cell visibility to cytotoxic CD8^+^ T cells.

Ferroptosis-associated oxidative stress has been shown to promote the release of damage-associated molecular patterns and tumor-associated antigens, thereby increasing antigen availability and immune activation, which in turn facilitates CD8^+^ T-cell activation and IFN-γ secretion. IFN-γ reinforces MHC class I expression and antigen processing machinery, sensitizing tumor cells to immune-mediated killing^45-47^. While ferroptosis can therefore potentiate MHC class I–mediated antitumor immunity, excessive or uncontrolled ferroptotic signaling can paradoxically disrupt antigen processing and promote immune escape^47^. Notably, our approach does not employ a canonical ferroptosis inducer; rather, inhibition of the MTDH– SND1 axis appears to impose a controlled susceptibility to ferroptotic stress. This regulated state may favor immunogenic remodeling without compromising antigen presentation capacity. Supporting this hypothesis, genetic deletion of SND1 has been shown to enhance MHC class I antigen presentation and increase CD8^+^ T-cell infiltration^48^, with mechanistic evidence that the MTDH–SND1 complex suppresses antigen presentation by destabilizing TAP1 and TAP2 transcripts and impairing peptide loading onto MHC class I–β2-microglobulin complexes^16^. Disruption of heavy chain–β2-microglobulin assembly in the endoplasmic reticulum impairs MHC class I antigen presentation, reducing tumor cell susceptibility to immune surveillance by cytotoxic CD8^+^ T cells. Downregulation of TAP proteins, particularly TAP1 and TAP2, represents a key mechanism of tumor immune evasion by limiting peptide loading onto MHC class I–β2m complexes and subsequent antigen presentation at the cell surface^49^. In line with these observations, we detected coordinated upregulation of TAP1, TAP2, and β2-microglobulin following combination treatment with olaparib and the MTDH–SND1 inhibitor C26-A6, compared with either monotherapy, in a BRCA-deficient model. Together, these findings support a model in which combination therapy modulates tumor antigen presentation through inhibition of the MTDH–SND1 axis, with potential consequences for adaptive immune recognition. Future studies will be required to define how this immunogenic remodeling influences therapeutic responses across diverse tumor contexts.

In conclusion, our study highlights the role of the MTDH–SND1 axis in shaping tumor cell stress responses and immunogenic features in BRCA-deficient ovarian cancer, providing a rationale for exploring combination strategies that extend beyond DNA repair–targeted therapies.

## Material and methods

### Cell lines and cell culture

Human ovarian cancer cell line PEO1 was cultured under conditions designed to closely mimic their physiological environment. Cells were maintained in RPMI-1640 medium supplemented with GlutaMAX™ (Gibco), 10% fetal bovine serum (FBS), and 1% penicillin-streptomycin (P/S). Mouse ovarian cancer cell lines PPNM and BPPNM were cultured in Dulbecco’s Modified Eagle Medium (DMEM) supplemented with 1% Insulin-Transferrin-Selenium (ITS-G, Gibco, #41400045), epidermal growth factor (EGF, 10 μg/mL), 4% FBS, and 1% P/S. All cell cultures were incubated at 37°C in a humidified atmosphere containing 5% CO_2_. Cells were passaged upon reaching approximately 80–90% confluency to maintain optimal growth conditions. All cell lines were tested for mycoplasma contamination prior to use and maintained under standard sterile culture conditions.

### Cell viability assay

Cell viability was assessed using the CellTiter-Glo Luminescent Cell Viability Assay (Promega, #G9242). Cells were seeded in 96-well plates (Corning) at a density of 300 cells for mousecell lines and 1500 cells per well for PEO1 and incubated overnight to allow attachment. Following drug treatment, cells were incubated for 5 days. At the end of the treatment period, CellTiter-Glo reagent was added directly to each well, followed by a 10-minute incubation in the dark. Luminescence, which is proportional to intracellular ATP levels and reflects the number of viable cells, was measured using a luminometer. For the rescue assay, cells were treated with erastin and/or C26-A6 in the presence or absence of ferrostatin-1 (2.5 µM). Ferrostatin-1 was added concurrently with treatment. Cell viability was assessed after 16 h using the CellTiter-Glo as described above. All experiments were performed in triplicate, and results were expressed as mean ± standard deviation (SD). The compounds used in this assay were erastin (Fisher Scientific, GTFHY-15763), ML-162 (Nordic Biosite, -20455), C26-A6 (Bio-Techne, -7692) and ferrostatin-1 (Fer-1; HY-100579; MedChem Express).

### Colony formation assay

Cells were seeded in 12-well plates at a density of 300 cells for mouse cell lines and 4000 cells per well for PEO1 and allowed to adhere overnight. Cells were then treated with compounds for 3 days, followed by incubation in fresh medium until visible colonies formed in the untreated control wells (approximately 10 days in total). Colonies were fixed and stained with a solution containing 6.0% glutaraldehyde and 0.5% crystal violet. Plates were air-dried at room temperature prior to imaging.

### Bliss synergy and IC_50_ analysis

To assess IC_50_ values and drug combination synergy, eight concentrations of each drug were tested individually and in combination, with all conditions performed in triplicate. Dose-response data were analyzed using DECREASE^50^, a machine learning–based method for predicting drug combination landscapes from limited experimental input. Experimental validation was performed using the CellTiter-Glo assay. IC_50_ values and synergy scores were calculated using the Bliss independence model via SynergyFinder^17^, a web-based tool for drug combination analysis. The Bliss model assumes that two drugs act independently, and their combined effect is estimated as the probability of independent events. Bliss synergy scores greater than zero indicate a possible synergistic interaction.

### Animal experiments

C57BL/6 female mice (8–12 weeks old; Jackson Laboratory, stock #000664) were used for *in vivo* studies. Prior to injection, three million cells were suspended in a 1:1 mixture of Matrigel (Corning, #47743-710) and FT medium. The cell suspension was administered intraperitoneally. Mice were treated intraperitoneally with olaparib (Selleckchem, S1060) at a dose of 50 mg/kg, twice per week^17^. For C26-A6 treatment, mice received 15 mg/kg, five times per week^15^. Both compounds were prepared for *in vivo* use according to the manufacturers’ instructions. Treatments were administered for a total duration of four weeks. Each experimental group consisted of 3–5 mice. Control mice received vehicle injections of DMSO at concentrations equivalent to those used for the corresponding treatment. Tumor samples were sectioned into pieces; one portion was immediately processed to isolate live cells for single-cell RNA sequencing, while another was snap-frozen in liquid nitrogen for subsequent proteomic and phosphoproteomic analyses. Abdominal area and tumor burden were quantified using Fiji (ImageJ, NIH), with image calibration performed using an in-frame ruler for accurate area measurement. All experiments were performed following approval by the Swedish Regional (Malmö–Lund) Ethical Committee (5.8.18-00866/2021) and in accordance with national and European Union guidelines. According to ethical permit individual tumor volume may not exceed 1 cm^3^. Tumor spread was assessed by measuring tumor area rather than volume. Although the largest tumor area observed in control mice was approximately 3 cm^2^, the estimated individual tumor volume remained well below 1 cm^3^.

### Protein extraction and digestion for proteomics

For proteomic analysis of cell lines, 4–5 biological replicates were processed per treatment and control condition. Before protein extraction, mouse and human cell lines were treated with C26-A6 and erastin as single agents or in combination (mouse cell lines: C26-A6 (65 μM), erastin (100 nM); human cell line: C26-A6 (95 μM), erastin (300 nM) for 72 h. Cells (at 70– 80% confluency) or tissues were lysed using boiling lysis buffer composed of 5% SDS (Thermo Fisher Scientific, Vilnius, Lithuania), 0.1 M Tris-HCl pH 8.5 (Thermo Scientific, Rockford, IL), 10 mM 2-chloroacetamide (2-CAA; Sigma-Aldrich, St. Louis, MO), and 5 mM TCEP (Thermo Scientific, Rockford, IL) prepared in LC/MS-grade water (Fisher Chemical, Ottawa, Canada), followed by sonication. Protein concentrations were estimated using the BCA assay, and 500 μg of protein was used for digestion. Protein aggregation capture (PAC)^51^ digestion was performed using a combination of LysC (1:500) and trypsin (1:250). Digestion was automated using a KingFisher Flex Robot (Thermo Fisher Scientific) with MagResyn Hydroxyl beads (Resyn Biosciences) in a 96-well format. Following digestion, peptides were desalted using 50 mg Sep-Pak tC18 Vac Cartridge columns (Waters, Milford, MA).

### Phosphopeptide enrichment

Phosphopeptide enrichment was performed using the KingFisher™ Flex Robot with MagResyn Zr-IMAC HP beads (Resyn Biosciences) in 96-well plates. Beads were first equilibrated in binding solvent consisting of 80% acetonitrile (MeCN), 5% trifluoroacetic acid (TFA), and 0.1 M glycolic acid (Sigma-Aldrich). Peptides were initially dissolved in 0.1% TFA at 4 μg/μL, then diluted in binding solvent and incubated with the equilibrated beads. After binding, the beads were washed sequentially with binding solvent, followed by 80% MeCN + 1% TFA, and 10% MeCN + 0.2% TFA. Phosphopeptides were eluted using 1% ammonia (pH ∼10) and immediately acidified with TFA to a final concentration of 1%.

### LC-MS data acquisition

Peptides and phosphopeptides were analyzed using an Evosep One LC system (Evosep) coupled to a Q Exactive HF-X Orbitrap mass spectrometer (Thermo Scientific). A total of 600 ng of peptides or the entire volume of enriched phosphopeptides, reconstituted in 20 μL of 0.1% formic acid, was loaded onto conditioned Evotips (Evosep). Chromatographic separation was performed on an in-house packed PicoFrit column using the 58-minute Whisper method (20 samples per day, SPD) for full proteome samples and the 32-minute Whisper Zoom method (40 SPD) for phosphopeptides. Mass spectrometry was conducted in data-independent acquisition (DIA) mode. For the 58-minute runs, MS1 scans ranged from 395–1005 m/z at a resolution of 60000, with a 55 ms maximum injection time and a 3e6 Automatic Gain Control (AGC) target. MS/MS were collected at 15000 resolution, 1e6 AGC target with automatic maximum injection time, NCE 27 with default charge 3, loop count of 75, and 151 staggered 8 m/z windows between 400.4 and 1000.7 m/z. For the 32-minute runs, MS1 scans covered 350–1400 m/z at 120000 resolution, with a 25 ms maximum injection time and a 3e6 AGC target. MS/MS scans were acquired at 15000 resolution, automatic injection time, 3e6 AGC target, NCE 27 with default charge 2, loop count of 50, and 100 staggered 13.7 m/z windows between 471.5 and 1149.7 m/z. All spectra for both methods were acquired in centroid mode.

### Data processing

Mass spectrometry data were processed in DIA-NN (v2.0 or 2.1.0)^52^. Full proteome RAW files were first demultiplexed and converted to mzML, while phospoenriched RAW files were used directly without prior file conversion. All analyses were performed in library-free mode to generate quantitative peptide- and protein-level data matrices. Human cell line and mouse model datasets were analyzed separately and searched against the corresponding reviewed UniProtKB proteomes (human proteome, 7 February 2025 version, 20504 proteins; mouse proteome, 6 December2024 version, 17334 proteins). Searches were conducted with tryptic specificity, allowing up to one missed cleavage. For proteome analyses, cysteine carbamidomethylation was specified as a fixed modification, and methionine oxidation, protein N-terminal methionine excision and protein N-terminal acetylation were included as a variable modifications with maximum 1 variable modification, except for the human cell line data were no variable modifications were set. For phosphoproteomic analyses, cysteine carbamidomethylation was specified as a fixed modification, while variable modifications included phosphorylation of serine, threonine, and tyrosine residues, protein N-terminal methionine excision and protein N-terminal acetylation. Phospho peptidoforms were inferred during database searching. Peptide- and protein-level identifications were filtered to a false discovery rate (FDR) of 1% within DIA-NN. The DIA-NN protein-level reports were used for further full proteome analyses, while the 90% phosphosite reports were used for phosphopeptide analysis.

### Data analysis

Data analysis was performed in R, (version 4.5.0). Normalization of protein- and phosphopeptide-level abundance data and differential expression analysis were performed using NormalyzerDE (v1.20)^53^, applying LIMMA empirical Bayes (eBayes) statistics. Multiple-testing correction was carried out using the Benjamini–Hochberg procedure, and group-wise differential expression was assessed at the specified false discovery rate (FDR) thresholds. OmicLoupe^54^ was used for the identification and visualization of differentially expressed proteins, including box plot representations.

Functional enrichment analyses were conducted using STRING^25^ and GSEA^21^, implemented with clusterProfiler^55^, to identify significantly enriched pathways and biological processes. Phosphorylation site–specific signaling changes were further assessed by PTM signature enrichment analysis (PTM-SEA)^26^ using PTMSigDB human flanking sequence signatures (version 2.0).

### Single-cell RNA seq

Fresh tissue samples were finely minced and digested (in digestion buffer containing 0.5 mg/mL collagenase, 1 mg/mL hyaluronidase, and 0.5 mg/mL DNase I in HBSS). Samples were incubated in a shaking incubator for 40 minutes. The resulting cell suspension was filtered through a 100 µm nylon cell strainer and centrifuged to discard the supernatant. The cell pellet was resuspended and passed through a 40 µm nylon cell strainer.

To remove erythrocytes, a 1× RBC lysis solution (Miltenyi Biotec, #130-094-183) was added to the cell suspension at a 9:1 ratio, followed by vortexing. Samples were incubated at room temperature for 2 minutes, then centrifuged at 300 × g for 10 minutes, and the supernatant was carefully aspirated. Dead cells were removed using the kit (Miltenyi Biotec, #130-090-101) according to the manufacturer’s instructions. Briefly, the cell pellet was resuspended in 100 µL of magnetic microbeads and diluted with 1× binding buffer. Magnetic columns were used for separation. Viable cells were counted using trypan blue exclusion staining. Single-cell suspensions were submitted to an external core facility for library preparation and sequencing using the 10x Genomics Chromium platform.

### Single-cell RNA sequencing data analysis

Raw sequencing data were processed using Cell Ranger (version 9.0.1,10x Genomics)^56^, and the resulting output files were imported into RStudio (version 4.3.2) following an initial quality control assessment based on Cell Ranger summary reports. Ambient RNA contamination was estimated and corrected at the single-cell level using SoupX (version 1.6.2)^57^.

Subsequent quality control and preprocessing were performed using Seurat (version 5.3.0)^58^. Cells with a total feature count below the 0.01 quantile were excluded. Cells were further filtered if the percentage of mitochondrial gene expression exceeded three standard deviations above the median and/or if the total UMI count was more than three standard deviations below the median. Mitochondrial genes were removed prior to downstream analyses.

Data normalization and variance stabilization were performed using SCTransform() (with the same median total UMI count for all samples), after which cell cycle scores were calculated using CellCycleScoring(), assigning S and G2/M phase scores to each cell, with G1 defined as the default phase. Dimensionality reduction was conducted by computing 100 principal components using PCA, and cells were visualized using Uniform Manifold Approximation and Projection (UMAP) based on the top 15 PCs.

Cell type annotation was initially performed using CellTypist (version 1.6.3), using a previously published murine scRNA-seq dataset (GEO accession GSM4813906) as reference^17,59^. The reference dataset was processed in Seurat following a comparable workflow, and unsupervised clustering was performed using the Louvain algorithm at a resolution of 0.1, yielding nine clusters. Cell type identities were assigned based on canonical marker gene expression as described in the original study^17^. A CellTypist model was trained on this annotated reference dataset and applied to each sample independently.

The normalized data of all samples were then merged using anndata (version 0.2.0) and further processed together using Scanpy (version 1.10.4). Highly variable genes (n = 2,000) were selected for downstream analysis. Dimensionality reduction was performed by principal component analysis (PCA). Batch effects were corrected using Harmony (harmonypy v0.0.10) based on the top 30 principal components. A k-nearest-neighbor graph was constructed, and cells were embedded in two dimensions using Uniform Manifold Approximation and Projection (UMAP) for visualization. Cells were grouped into distinct clusters based on similar gene expression profiles, potentially reflecting their biological identities, using graph-based clustering via the Leiden algorithm with a resolution parameter set to 0.2. Leveraging automatic cell type annotations generated by CellTypist, we consolidated several clusters representing fine-grained cellular heterogeneity, resulting in a final set of nine distinct cell clusters. We then identified cluster-specific marker genes by performing differential expression analysis using Wilcoxon rank-sum tests. The biological identity of each cluster was manually assigned by integrating automatic cell type annotations generated by CellTypist, cluster-specific marker genes, and established marker genes corresponding to expected cell types.

### Proteomics cell-type deconvolution

Cell-type composition of mouse tumors was estimated from bulk tumor proteomics using proteoDeconv (running the official CIBERSORTx Docker image)^60,61^. Cell-type marker genes were derived from the tumor scRNA-seq dataset generated in this study and used to assign cell-type labels in an external scRNA-seq dataset (GEO GSE158474)^17^. The labeled GSE158474 expression matrix was exported in CIBERSORTx single-cell reference format (27,998 genes) and used for CIBERSORTx single-cell signature matrix construction (replicates=5, q.value=0.01, G.min=300, G.max=500).

For deconvolution, protein-group intensities were collapsed to gene-level abundances (first symbol when multiple were present). Missing values were imputed to the lowest observed non-zero value across the matrix, duplicated gene symbols were collapsed by retaining the entry with the highest median abundance, and each sample was converted to a TPM-like relative-abundance matrix (scaled to 1e6 per sample). Deconvolution was then performed with CIBERSORTx^60^ via proteoDeconv^61^ to obtain relative cell-type fractions per sample. Group differences were assessed per cell type using two-sided Wilcoxon rank-sum tests without multiple-testing correction.

## Data availability

The mass spectrometry proteomics data have been deposited to the ProteomeXchange Consortium via the PRIDE^62^ partner repository with the dataset identifier PXD073933.

## Acknowledgements

This study was funded by Fru Berta Kamprads Stiftelse FBKS 2021-31(330), Mats Paulssons Stiftelse, Stiftelsen Cancera and the technical faculty at Lund University through Proteoforms@LU. Some computations were enabled by resources provided by the National Academic Infrastructure for Supercomputing in Sweden (NAISS, project 2025/22-62), partially funded by the Swedish Research Council through grant agreement no. 2022-06725. Mouse ovarian cancer cell lines PPNM and BPPNM were kindly provided by the laboratory of Professor Robert A. Weinberg^17^.

## Contributions

PE, AN, EP, and BB conducted experiments. PE, EE, MZ and FL analyzed the data. PE, AN, JKU, and FL developed the methodology and contributed to the study design. ASG, JKU, YL and FL supervised the work. FL acquired funding. PE drafted the manuscript. All authors edited and approved the final manuscript.

## References

1 Wang, Y., Duval, A. J., Adli, M. & Matei, D. Biology-driven therapy advances in high-grade serous ovarian cancer. The Journal of Clinical Investigation 134 (2024).

2 Lheureux, S., Gourley, C., Vergote, I. & Oza, A. M. Epithelial ovarian cancer. The Lancet 393, 1240–1253 (2019).

3 Toss, A. et al. Hereditary ovarian cancer: not only BRCA 1 and 2 genes. BioMed research international 2015, 341723 (2015).

4 Hinchcliff, E. M., Bednar, E. M., Lu, K. H. & Rauh-Hain, J. A. Disparities in gynecologic cancer genetics evaluation. Gynecologic oncology 153, 184–191 (2019).

5 Ren, N. et al. Efficacy and safety of PARP inhibitor combination therapy in recurrent ovarian cancer: a systematic review and meta-analysis. Frontiers in Oncology 11, 638295 (2021).

6 Sánchez-Lorenzo, L., Salas-Benito, D., Villamayor, J., Patiño-García, A. & González-Martín, A. The BRCA gene in epithelial ovarian cancer. Cancers 14, 1235 (2022).

7 Dinkins, K. et al. Targeted therapy in high grade serous ovarian Cancer: A literature review. Gynecologic Oncology Reports, 101450 (2024).

8 Konstantinopoulos, P. A., Ceccaldi, R., Shapiro, G. I. & D’Andrea, A. D. Homologous recombination deficiency: exploiting the fundamental vulnerability of ovarian cancer. Cancer discovery 5, 1137–1154 (2015).

9 Lei, G. et al. BRCA1-mediated dual regulation of ferroptosis exposes a vulnerability to GPX4 and PARP co-inhibition in BRCA1-deficient cancers. Cancer discovery 14, 1476–1495 (2024).

10 Xie, X. et al. Targeting GPX4-mediated ferroptosis protection sensitizes BRCA1-deficient cancer cells to PARP inhibitors. Redox Biology 76, 103350 (2024).

11 Ojo, O. A. et al. Ferroptosis as the new approach to Cancer therapy. Cancer Treatment and Research Communications, 100913 (2025).

12 Zhou, Q. et al. Ferroptosis in cancer: from molecular mechanisms to therapeutic strategies. Signal transduction and targeted therapy 9, 55 (2024).

13 Esmaeili, P. et al. Proteomics discovery of MTDH and SND1 interaction vulnerabilities in ovarian cancer. Scientific Reports (2025).

14 Blanco, M. A. et al. Identification of staphylococcal nuclease domain-containing 1 (SND1) as a Metadherin-interacting protein with metastasis-promoting functions. Journal of Biological Chemistry 286, 19982–19992 (2011).

15 Shen, M. et al. Small-molecule inhibitors that disrupt the MTDH–SND1 complex suppress breast cancer progression and metastasis. Nature cancer 3, 43–59 (2022).

16 Shen, M. et al. Pharmacological disruption of the MTDH–SND1 complex enhances tumor antigen presentation and synergizes with anti-PD-1 therapy in metastatic breast cancer. Nature cancer 3, 60–74 (2022).

17 Iyer, S. et al. Genetically defined syngeneic mouse models of ovarian cancer as tools for the discovery of combination immunotherapy. Cancer Discovery 11, 384–407 (2021).

18 Chen, Z. et al. Ferroptosis as a potential target for cancer therapy. Cell death & disease 14, 460 (2023).

19 Bi, J. et al. Metadherin enhances vulnerability of cancer cells to ferroptosis. Cell death & disease 10, 682 (2019).

20 Ianevski, A., Giri, A. K. & Aittokallio, T. SynergyFinder 3.0: an interactive analysis and consensus interpretation of multi-drug synergies across multiple samples. Nucleic acids research 50, W739–W743 (2022).

21 Subramanian, A. et al. Gene set enrichment analysis: a knowledge-based approach for interpreting genome-wide expression profiles. Proceedings of the National Academy of Sciences 102, 15545–15550 (2005).

22 Tang, D., Chen, X., Kang, R. & Kroemer, G. Ferroptosis: molecular mechanisms and health implications. Cell research 31, 107–125 (2021).

23 Yan, R. et al. NRF2, a superstar of ferroptosis. Antioxidants 12, 1739 (2023).

24 Kolberg, L. et al. g: Profiler—interoperable web service for functional enrichment analysis and gene identifier mapping (2023 update). Nucleic acids research 51, W207–W212 (2023).

25 Szklarczyk, D. et al. The STRING database in 2021: customizable protein–protein networks, and functional characterization of user-uploaded gene/measurement sets. Nucleic acids research 49, D605–D612 (2021).

26 Krug, K. et al. A curated resource for phosphosite-specific signature analysis. Molecular & cellular proteomics 18, 576–593 (2019).

27 Wan, L. et al. MTDH-SND1 interaction is crucial for expansion and activity of tumor-initiating cells in diverse oncogene-and carcinogen-induced mammary tumors. Cancer cell 26, 92–105 (2014).

28 Shen, H. et al. Overcoming MTDH and MTDH-SND1 complex: Driver and potential therapeutic target of cancer. Cancer Insight 3, 55–82 (2023).

29 Kim, J. W., Lee, J.-Y., Oh, M. & Lee, E.-W. An integrated view of lipid metabolism in ferroptosis revisited via lipidomic analysis. Experimental & Molecular Medicine 55, 1620–1631 (2023).

30 Ding, X. et al. Ferroptosis in cancer: revealing the multifaceted functions of mitochondria. Cellular and Molecular Life Sciences 82, 277 (2025).

31 Cañeque, T. et al. Activation of lysosomal iron triggers ferroptosis in cancer. Nature, 1–9 (2025).

32 Lecona, E. & Fernandez-Capetillo, O. Targeting ATR in cancer. Nature Reviews Cancer 18, 586–595 (2018).

33 Gudmundsdottir, K. & Ashworth, A. The roles of BRCA1 and BRCA2 and associated proteins in the maintenance of genomic stability. Oncogene 25, 5864–5874 (2006).

34 Li, X. & Zou, L. BRCAness, DNA gaps, and gain and loss of PARP inhibitor–induced synthetic lethality. The Journal of clinical investigation 134 (2024).

35 Haynes, B., Murai, J. & Lee, J.-M. Restored replication fork stabilization, a mechanism of PARP inhibitor resistance, can be overcome by cell cycle checkpoint inhibition. Cancer treatment reviews 71, 1–7 (2018).

36 Bhamidipati, D., Haro-Silerio, J. I., Yap, T. A. & Ngoi, N. PARP inhibitors: enhancing efficacy through rational combinations. British journal of cancer 129, 904–916 (2023).

37 Hong, T. et al. PARP inhibition promotes ferroptosis via repressing SLC7A11 and synergizes with ferroptosis inducers in BRCA-proficient ovarian cancer. Redox biology 42, 101928 (2021).

38 Tew, W. P. et al. PARP inhibitors in the management of ovarian cancer: ASCO guideline. Journal of Clinical Oncology 38, 3468–3493 (2020).

39 Xiao, Y. et al. Emerging therapies in cancer metabolism. Cell metabolism 35, 1283–1303 (2023).

40 Surace, L. et al. Complement is a central mediator of radiotherapy-induced tumor-specific immunity and clinical response. Immunity 42, 767–777 (2015).

41 Lee, K. S., Zhang, Q., Suwa, T., Clark, H. & Olcina, M. M. In Seminars in Immunology. 101927 (Elsevier).

42 Dhillon, A. S., Hagan, S., Rath, O. & Kolch, W. MAP kinase signalling pathways in cancer. Oncogene 26, 3279–3290 (2007).

43 Conte, A., Valente, V., Paladino, S. & Pierantoni, G. M. HIPK2 in cancer biology and therapy: Recent findings and future perspectives. Cellular Signalling 101, 110491 (2023).

44 Kwon, J., Zhang, J., Mok, B. & Han, C. CK2-mediated phosphorylation upregulates the stability of USP13 and promotes ovarian cancer cell proliferation. Cancers 15, 200 (2022).

45 Qi, D. & Peng, M. Ferroptosis-mediated immune responses in cancer. Frontiers in Immunology 14, 1188365 (2023).

46 Wahida, A. & Conrad, M. Decoding ferroptosis for cancer therapy. Nature Reviews Cancer, 1–15 (2025).

47 Zhao, X., Li, X. & Xu, Y. Ferroptosis: a dual-edged sword in tumour growth. Frontiers in pharmacology 14, 1330910 (2024).

48 Wang, Y. et al. Oncoprotein SND1 hijacks nascent MHC-I heavy chain to ER-associated degradation, leading to impaired CD8+ T cell response in tumor. Science Advances 6, eaba5412 (2020).

49 Rai, S. K. et al. Staphylococcal nuclease and tudor domain-containing protein 1: An emerging therapeutic target in cancer. Molecular and Clinical Oncology 23, 86 (2025).

50 Ianevski, A. et al. Prediction of drug combination effects with a minimal set of experiments. Nature machine intelligence 1, 568–577 (2019).

51 Batth, T. S. et al. Protein aggregation capture on microparticles enables multipurpose proteomics sample preparation. Molecular & Cellular Proteomics 18, 1027–1035 (2019).

52 Demichev, V., Messner, C. B., Vernardis, S. I., Lilley, K. S. & Ralser, M. DIA-NN: neural networks and interference correction enable deep proteome coverage in high throughput. Nature methods 17, 41–44 (2020).

53 Willforss, J., Chawade, A. & Levander, F. NormalyzerDE: online tool for improved normalization of omics expression data and high-sensitivity differential expression analysis. Journal of proteome research 18, 732–740 (2018).

54 Willforss, J., Siino, V. & Levander, F. OmicLoupe: facilitating biological discovery by interactive exploration of multiple omic datasets and statistical comparisons. BMC bioinformatics 22, 107 (2021).

55 Yu, G. Thirteen years of clusterProfiler. The Innovation 5 (2024).

56 Zheng, G. X. et al. Massively parallel digital transcriptional profiling of single cells. Nature communications 8, 14049 (2017).

57 Young, M. D. & Behjati, S. SoupX removes ambient RNA contamination from droplet-based single-cell RNA sequencing data. Gigascience 9, giaa151 (2020).

58 Hao, Y. et al. Dictionary learning for integrative, multimodal and scalable single-cell analysis. Nature biotechnology 42, 293–304 (2024).

59 Domínguez Conde, C. et al. Cross-tissue immune cell analysis reveals tissue-specific features in humans. Science 376, eabl5197 (2022).

60 Newman, A. M. et al. Determining cell type abundance and expression from bulk tissues with digital cytometry. Nature biotechnology 37, 773–782 (2019).

61 Zamore, M. n. et al. Considerations and software for successful immune cell deconvolution using proteomics data. Journal of Proteome Research 24, 3751–3761 (2025).

62 Perez-Riverol, Y. et al. The PRIDE database at 20 years: 2025 update. Nucleic acids research 53, D543–D553 (2025).

