## Supplemental information for "MTDH-SND1 disruption sensitizes ovarian cancer to ferroptosis and PARP inhibition"

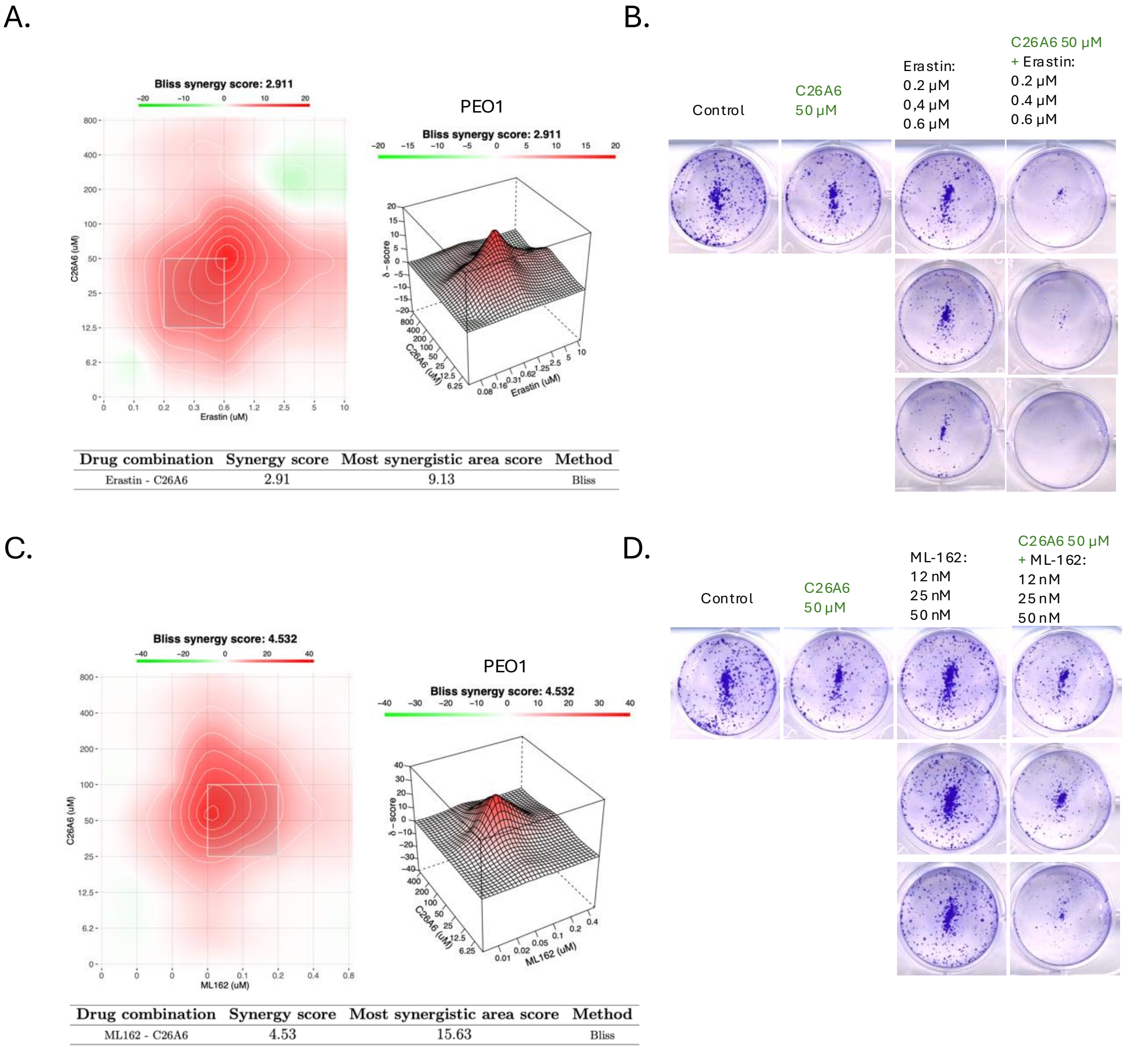


**Supplementary Figure 1. Synergistic effect of C26A6 with Erastin or ML-162 in PEO1 cells.**
**A.** Drug interaction analysis of C26-A6 in combination with erastin
**B.** Colony formation assay in PEO1 cells treated with vehicle, C26-A6 (50 µM), erastin (0.4 or 0.6 µM), or their combination. Combination treatment reduced colony number and size.
**C.** Drug interaction analysis of C26-A6 in combination with ML-162.
**D.** Colony formation assay in PEO1 cells treated with vehicle, C26-A6 (50 µM), ML-162 (12 or 25 nM), or their combination. A marked reduction in colony number and size was observed with the combination treatment.

A.

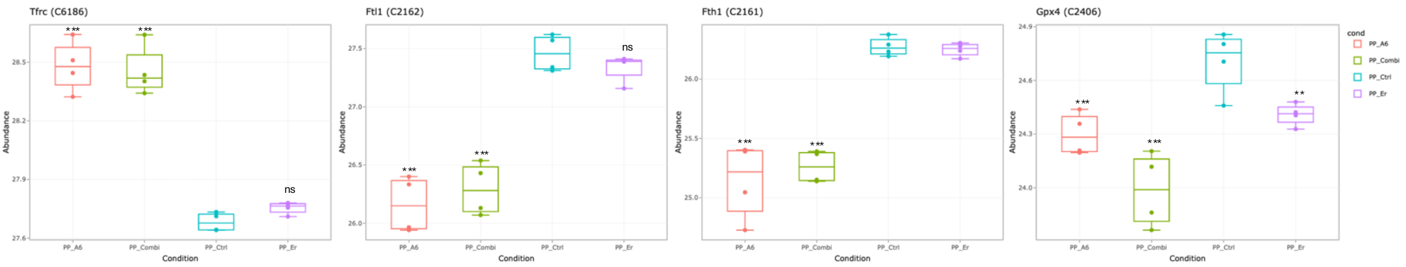


B.

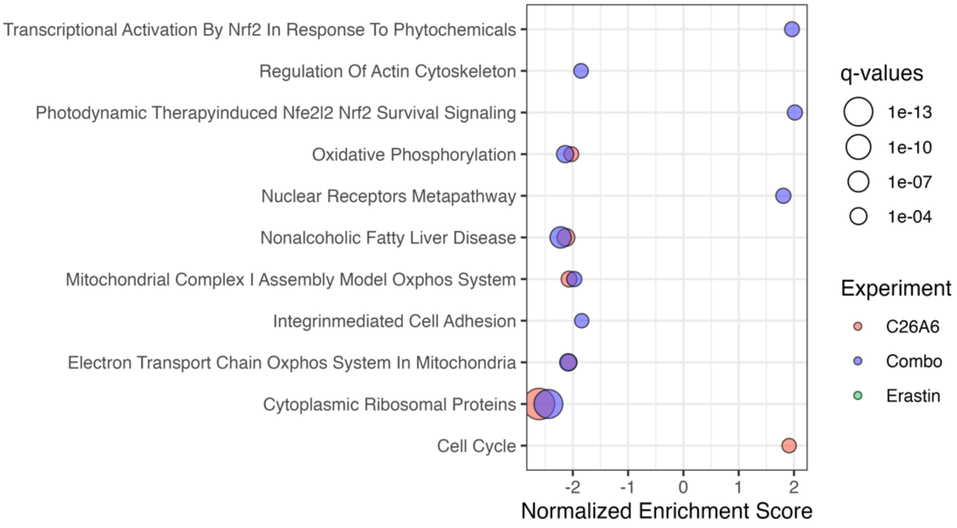


**Supplementary Figure 2. Proteomic profiling in PPNM**

**A.** Box plots showing the abundance of TFRC, FTL1, FTH1, and GPX4 protein expression in PPNM cells under erastin, C26-A6, combination, and untreated conditions.

Statistical significance is indicated as follows: ns, not significant, $p < 0.05$ (*), $p < 0.01$ (**), $p < 0.001$ (***). All comparisons are relative to the control group. All comparisons are relative to the control group.

**B.** GSEA of PPNM across all treatment conditions, with pathways shown at a significance threshold of pvalueCutoff < 0.01.

A.

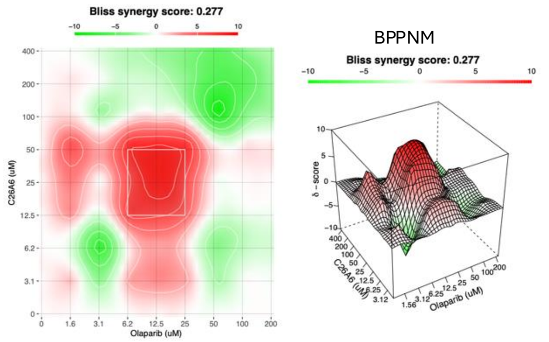


| Drug combination | Synergy score | Most synergistic area score | Method |
|---|---|---|---|
| Olaparib - C26A6 | 0.28 | 6.93 | Bliss |

B.

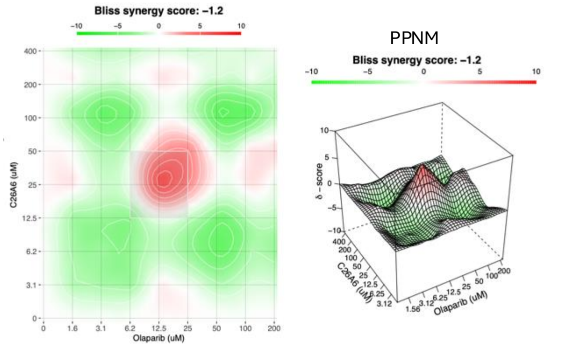


| Drug combination | Synergy score | Most synergistic area score | Method |
|---|---|---|---|
| Olaparib - C26A6 | -1.20 | 0.93 | Bliss |

C.

BPPNM

Control C26A6 200 uM Olaparib 6 uM Combination

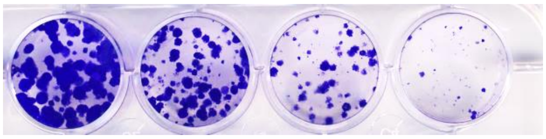

**Supplementary Figure 3. Disruption of the MTDH–SND1 interaction enhances sensitivity to PARP inhibition in BRCA1-deficient cells.**

**A.** Bliss synergy analysis of combined C26-A6 and olaparib treatment in BRCA1-deficient BPPNM cells, showing synergistic interaction (Bliss score = 6.93).

**B.** Bliss synergy analysis of the same drug combination in BRCA-proficient PPNM cells, showing minimal synergy (Bliss score = 0.93).

**C.** Clonogenic survival assay of BPPNM cells treated with C26-A6, olaparib, or the combination.

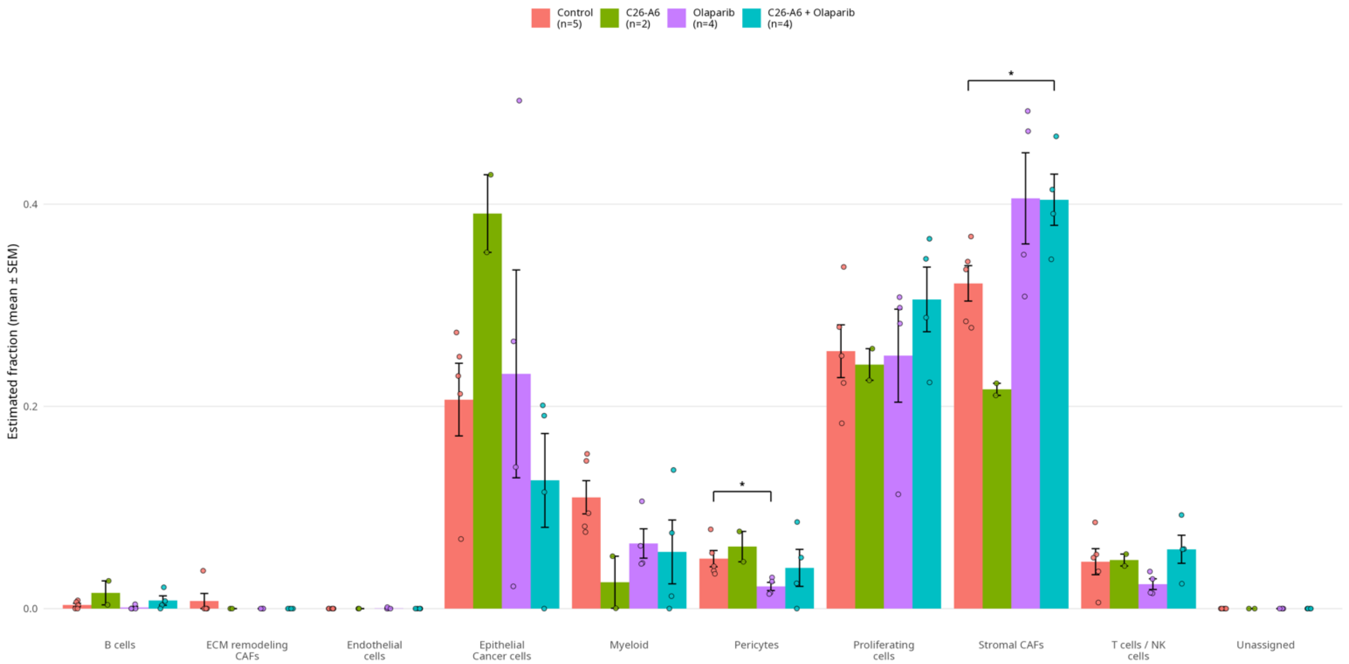


**Supplementary Figure 4.** Estimated cell-type fractions in bulk tumor proteomes inferred by CIBERSORTx via the proteoDeconv package using a custom single-cell–derived signature matrix based on the GSE158474 dataset. Bars show mean ± SEM and points show individual tumors per group (Control, C26-A6, C26-A6 + olaparib, olaparib). Pairwise group differences were assessed per cell type using two-sided Wilcoxon rank-sum tests without multiple-testing correction, only nominally significant comparisons ($p < 0.05$) are annotated.

**Supplementary Table 1.** Summary of the detected cells in scRNA-seq data.

| Cell type | Cell type distribution (%) | | | |
| --- | --- | --- | --- | --- |
| | Control | C26A6 | Olaparib | Combination |
| Stromal/CAFs | 64.92 | 55.43 | 59.6 | 64.11 |
| Proliferating cells | 16.48 | 24.61 | 33.24 | 22.52 |
| Epithelial / Cancer cells | 3.8 | 5.69 | 1.17 | 4.65 |
| Endothelial cells | 1.71 | 2.97 | 0.09 | 0.87 |
| B cells | 0.02 | 0.04 | 0.05 | 0.05 |
| Myeloid | 9.3 | 6.35 | 1.91 | 5.13 |
| ECM remodeling CAFs | 1.25 | 2.78 | 2.86 | 1.39 |
| T cells/NK cells | 2.07 | 1.51 | 0.96 | 1.07 |
| Pericytes | 0.45 | 0.61 | 0.11 | 0.21 |
